# Map simulator of tick abundance in heterogeneous agricultural landscapes

**DOI:** 10.1101/2025.07.08.663759

**Authors:** Gwenaël Vourc’h, David Abrial, Albert Agoulon, Karen D. McCoy, Alain Butet, Hélène Verheyden, Robin Loche\, Isabelle Lebert, Grégoire Perez, Elsa Quillery, Amélie Chastagner, Elsa Léger, Yann Rantier, A.J. Mark Hewison, Nicolas Morellet, Suzanne Bastian, Thierry Hoch, Olivier Plantard

**Affiliations:** Université Clermont Auvergne, INRAE, VetAgro Sup, UMR EPIA, 63222 Saint Genès Champanelle, France; Université Lyon, INRAE, VetAgro Sup, UMR EPIA, 69280 Marcy l’Etoile, France; Oniris, INRAE, BIOEPAR, 44300 Nantes, France; University of Montpellier CNRS IRD, UMR MIVEGEC, 34394 Montpellier, France; UMR CNRS 6553–Université de Rennes 1, ECOBIO, Campus de Beaulieu, 35042 Rennes, France; Université de Toulouse, INRAE, CEFS, 31326 Castanet-Tolosan, France; LTSER ZA PYrénées GARonne, 31320 Auzeville-Tolosane, France

**Keywords:** *Ixodes ricinus*, spatial distribution, statistical modeling, simulation, woodland ecotone, hedgerow

## Abstract

Among vector-borne diseases, tick-borne diseases (TBD) are a major concern for human health. Mapping the distribution of important tick species is thus a major challenge for efficient prevention. Due to its specific ecological requirements, *Ixodes ricinus*, the main tick species in Europe responsible for TBD transmission, lives mostly in woodlands but also at the interface between woodlands and pastures or crops and along hedgerows. At the landscape scale, extensive variations in tick densities are observed but remain poorly understood. In that aim, we built a statistical model to identify the landscape variables influencing the abundance of questing *I. ricinus* nymphs, using GLMM approaches and MCMC estimates. This model was fitted on a data set based on a field sampling of ticks conducted during 3 years in 2 different agricultural landscapes in northwest and southwest France, for a total of 5390 sampling units. Among 12 variables investigated, 4 were finally kept in the model: woodland perimeter, woodland distance, road distance and building perimeter. Then, we developed a *R* package that simulates the abundance of questing nymphs within a given agricultural landscape, taking into account the influence of the different habitats as determined by the above statistical model. The maps obtained as an output from this simulator will be a useful tool for visualizing TBD risk, notably for stake-holders involved in landscape management and public health decisions.

**Graphical abstract:** 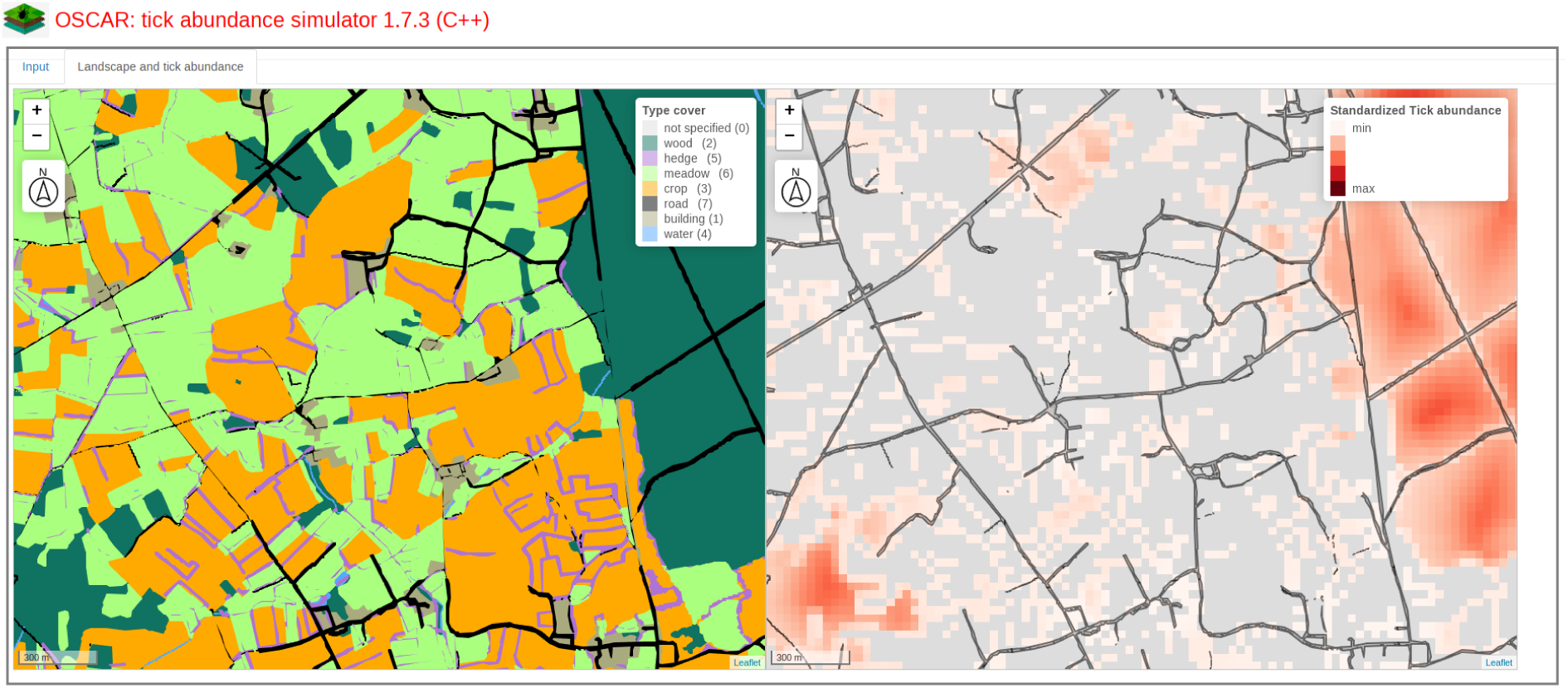

**Highlights:** *Ixodes ricinus* abondance is influenced by landscape characteristics

Tick sampling was carried out in heterogeneous agricultural landscapes

Informative variables related to habitats were identified by statistical analysis

Woodlands, roads and buildings influence tick densities

The resulting model was used to build a simulator of tick at-risk zones

## 1. Introduction

Determining the spatial distribution of parasites and pathogens, from the intra-host to the global scale, is essential for understanding and predicting the spread of diseases in space and time, and therefore for disease control (Anderson & May 1991). This knowledge is especially crucial for vector-borne diseases, where abiotic and biotic environmental factors vary across the spatial scales considered and have a considerable impact on their epidemiology through the distribution and activity of the vectors. Ticks are second only to mosquitoes in terms of their contribution to human vector-borne diseases worldwide, but are first if only temperate zones are considered (de la Fuente et al. 2008). Like most parasites, ticks have an aggregated distribution due to various factors, notably in the spatial aggregation of their hosts (Shaw & Dobson 1995). *Ixodes ricinus*, the main European vector of *Borrelia burgdorferi* sensu lato and Tick-Borne-Encephalitis virus, the etiologic agents of Lyme borreliosis and Tick-Borne encephalitis respectively, is widespread from Ireland to Turkey and from Spain to Scandinavia (Noll et al. 2023). However, on a finer scale, such as that of the landscape (Turner et al. 2001), *I. ricinus* densities vary widely: for the nymphal stage, the most important life stage from a human health perspective (Barbour & Fish 1993), densities estimated by the classical dragging method range from 0 to more than several hundreds of nymphs per 10 m^2^ (Boyard et al. 2007, Lebert et al. 2020, Vourc’h et al. 2016). As spatial heterogeneity in vector density induces spatial variation in the risk of tick-borne disease (Braks et al. 2016), understanding the complex links between landscape heterogeneity and tick distribution is a central challenge. Landscape heterogeneity can influence tick distribution in two ways. First, different habitat types are more or less favorable to tick survival. As free-living ticks are found on the ground or on herbaceous vegetation, they are influenced by temperature and humidity, two key drivers affecting tick physiology (Ogden et al. 2021). These abiotic conditions are also directly influenced by vegetation, including forest structure but also woody and herb layers (Vanroy et al. 2024). Second, the distribution of tick hosts - which participate in tick development, dispersal and pathogen transmission - is influenced by landscape heterogeneity. For example, hosts such as roe deer (*Capreolus capreolus*) - that host all life stages of *I. ricinus*, with a particular key host role for the adult female stage (Krol et al. 2020, Vor et al. 2010) - show marked preference for wooded habitats over more open ones, where they are exposed to hunting or other human disturbance (Kiffner et al. 2010, Martin et al. 2018). The distribution at the landscape-scale of small mammals such as various rodent and shrew species, major hosts of *I. ricinus* larvae and main reservoirs of several tick-borne pathogens, is also largely conditioned by habitat heterogeneity, which influences both food resources and predation risk (Fuentes-Montemayor et al. 2020). In addition, the size of habitat patches can influence the distribution of ticks. For example, the probability of local extinction of mammal host populations is higher in small patches of woodland, than in large forests due to greater fluctuations in population size and threshold effects (Guzzetta et al. 2017, Lloyd-Smith et al. 2005). As large herbivores and small mammals have very different home range sizes, these dynamics will vary depending on the host species.

In Europe, heterogeneous agricultural landscapes occupy vast areas and comprise a diversity of habitats including woodlands, hedgerows (which can act as linear corridors between wooded patches), crops and livestock pastures (Baudry et al. 2000, Litza et al. 2022). There is growing interest in accounting for the spatial heterogeneity of habitats at the landscape scale to predict variations in tick abundance in space (Diuk-Wasser et al. 2021, Estrada-Peña 2009, Gilbert 2016). Most studies are based on satellite data, enabling the construction of maps inferring tick density on a large scale, e.g. national or continental (see for example Asghar et al. 2016, Boehnke et al. 2015, Ehrmann et al. 2017, Estrada-Peña et al. 2016, Knoll et al. 2021, Li et al. 2016, Medlock et al. 2022, Qviller et al. 2013, 2016, Tack et al. 2012, Tran & Waller 2013, Zolnik et al. 2015). Nevertheless, despite their functional importance in these agroecosystems, hedgerows are often too small to be detected by this type of approach, and so are rarely taken into account in these studies (but see Boyard et al. 2008, Ehrmann et al. 2017, Estrada-Peña 2002, Medlock et al. 2020, Perez et al. 2016 or Perez et al. 2020).

Predicting risks related to ticks in agricultural landscapes requires thus to create risk maps that take into account habitat heterogeneity in these areas. To our knowledge, there is no existing simulator of tick abundance at the landscape scale. Beugnet et al. (2009) developed a tool to simulate the evolution of tick activity but at very large scales (*id est* a country), with a resolution of 27 km^2^. At a similar scale, Li et al. (2016) used a model to simulate the impact of climate change on Lyme disease risk and its seasonality in the heterogeneous landscapes of Scotland. At finer scales, Li et al. (2012) simulated the effect of landscape fragmentation on Lyme disease risk through a theoretical approach. A theoretical landscape was also built by Agudelo et al. (2021) to investigate the effect of white-tailed deer habitat use preferences on southern cattle fever tick. Wang et al. (2012) simulated the effect of the implantation of a green belt in an existing landscape on the abundance of *Amblyomma americanum*. Their agent-based model, which was not compared with observations, was however more dedicated to the test of such an hypothesis than to simulate tick-borne risk at a landscape scale. The aim of our study was to build a prototype simulator to map the estimated abundance of nymphs of *I. ricinus* on vegetation in agricultural landscapes, as a function of key drivers of spatial heterogeneity in the environment. This map simulator of nymphal abundance should be generic and usable on any similar landscape provided that the description of habitat types and their heterogeneity are available. To do so, we i) sampled *I. ricinus* nymphs on vegetation in two agricultural areas of France, ii) determined the statistical relationship between specific drivers of habitat heterogeneity and nymph abundance using GLMMs and Monte-Carlo Markov Chain (MCMC) estimates and iii) developed a *R* package that simulates nymph abundance within an agricultural landscape, taking into account the influence of those habitat features identified as informative by the statistical model.

## 2. Material and Methods

All the data analyzed in this article and a description of field collection methods are available in a data paper (Lebert et al. 2020). We therefore present only the information required for a general understanding of the methods here.

### 2.1. Study areas

The two study areas (Figure 1) are heterogeneous agricultural landscape composed of crops and natural meadows, interspersed with woodlands and hedgerows, both part of the International Long-Term Ecological Research (ILTER) network (“Zones Ateliers”, http://www.za-inee.org/en/node/804). The first area, the ‘Zone Armorique’ (‘ZAM’, now renamed ZAAr), is located in the North-West of France (N 48° 29’, E 1° 37’, https://osur.univ-rennes1.fr/zaar-home). Within its 147 km^2^ surface is a large forest of about 1000 ha (forests being considered as woodlands larger than 50 ha), many woods (woodlands smaller than 50 ha) and hedgerows (defined as a patch of woody vegetation with a maximal width of 5 m: Agoulon et al. 2012) surrounding meadows and crop fields.

**Figure 1:**
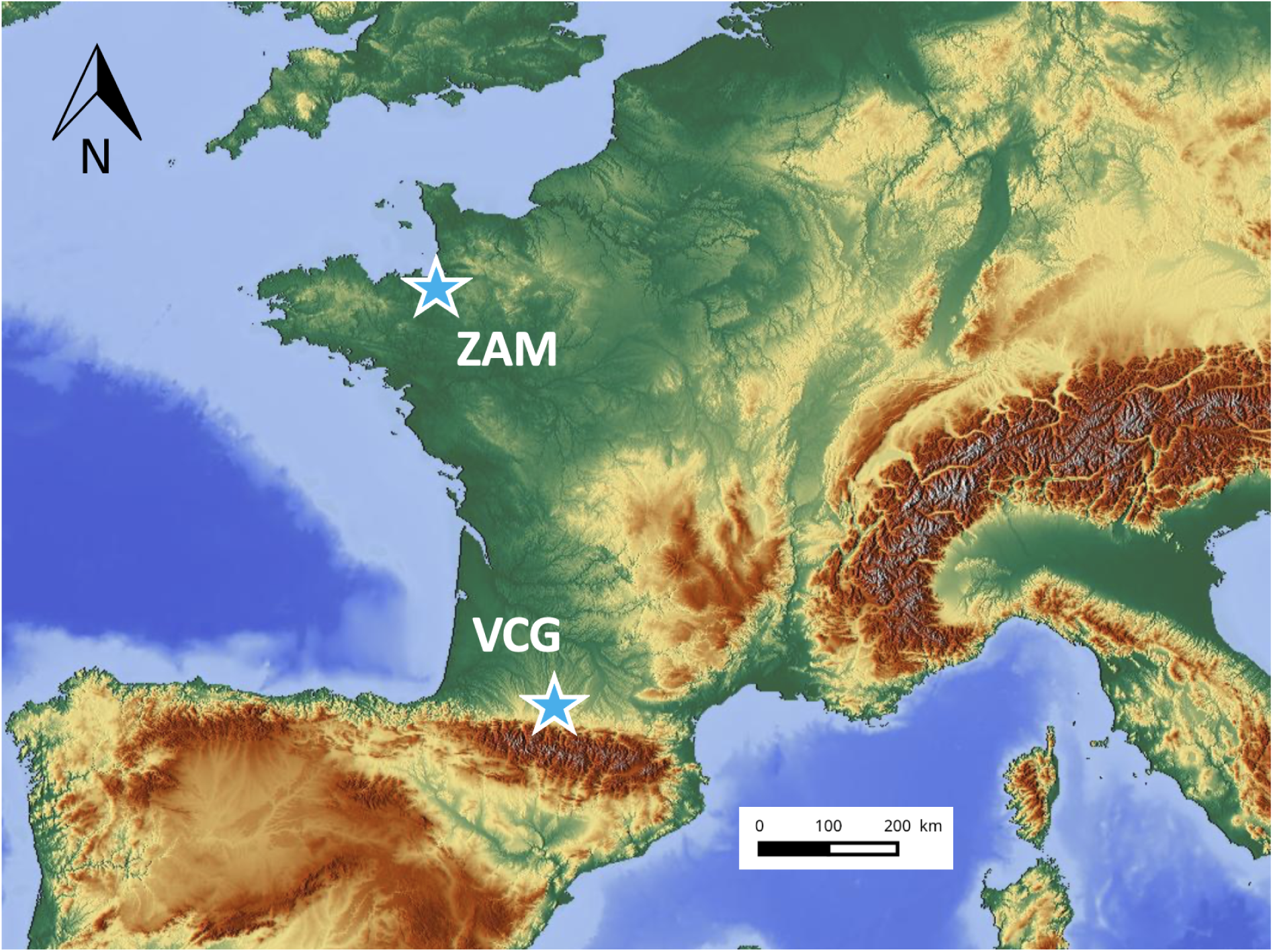
Localization of the two study areas, ‘Zone Armorique’ (ZAM) and ‘Vallées et Coteaux de Gascogne’ (VCG). Background map from https://maps-for-free.com/

The second area is located in the South-West of France (N 43° 15’, E 0° 51’) in the ‘Vallées et Coteaux de Gascogne’ (‘VCG’, now renamed PYGAR, https://pygar.omp.eu/). This 212 km^2^ area includes two large forest patches (∼500 and 700 ha) in addition to many woods and hedgerows.

### 2.2. Sampling of *Ixodes ricinus* nymphs on vegetation

In the springs of 2012 (May), 2013 (May) and 2014 (April), *I. ricinus* nymphs were collected on vegetation at ZAM and VCG areas by drag sampling (MacLeod 1932, Vassallo et al. 2000).

In each area, sampling zones (n = 60) were defined in 4 landscape types: agricultural landscapes with a low (1) or a high (2) hedgerow network density; forest edge (3); forest core (4) (Lebert et al. 2020).

In each sampling zone, one or two transect lines were defined: 1) a single transect line was sampled in the meadows along hedgerows (landscape types 1 or 2) and in the forest core (landscape type 4); 2) two transect lines were run along forest edges, on both sides of the ecotone (landscape type 3). This resulted in a total of 90 transect lines.

Each transect line included ten sampling units (10 m x 1 m = 10 m^2^), which were each separated by 20 m. Consequently, each transect line represented a sampling area of 100 m^2^. A total of 5,390 sampling units were surveyed over the three years (10 sampling units were missing due to the lack of a transect line in the forest core of ZAM in 2012). The same transect lines were visited each year.

In each sampling unit, a 1 m x 1 m white flannel cloth was slowly dragged (0.5 m/s) along 10 m across the low vegetation or leaf-litter (Agoulon et al. 2012, Perez et al. 2016). Collected ticks were immediately stored in 70% ethanol. Ticks were later identified in the laboratory using morphological criteria with a binocular microscope (Pérez-Eid 2007).

### 2.3. Explanatory variables considered in the statistical model to explain the abundance of *Ixodes ricinus* nymphs

We first selected meteorological and landscape variables (see Table 1) that could be measured in any location (notably remote-measured variables, obtained by satellites). We avoided fine scale field-measured variables, such as the detailed composition of vegetation cover in each sampling unit, to enable a wide applicability of this methodological approach. Among remote-measured variables, we selected those potentially influencing nymphal abundance (Vourc’h et al. 2016). In particular, we considered the presence of hedgerows based on the human photo-interpretation of satellite data. We also included the time of sampling (week and time of day) to account for a potential effect of within- and between-day variation during the sampling period. In addition, we included variables that were specifically linked to our field campaigns (area, year of sampling) as random effects, to control for repeated observations per area and year (Kendall and Stuart 1968) (see the following part on statistical analysis). The considered meteorological data were temperature (TEM) and humidity (HUM). These were obtained from ‘Météo France’ (https://donneespubliques.meteofrance.fr), which provided hourly data from 2011 to 2014. These meteorological variables were averaged over a determined period of time. The length of the temporal window considered to calculate the mean value was assessed using a Pearson correlation between nymphal abundance on a particular date and each meteorological variable, increasing the time step by 1 hour and including up to 30 days prior to sampling (the start time, “0”, was the hour of sampling). Thus, the best period for averaging was determined to be the previous 5 days before sampling (0-120 h) for TEM and the previous 14 days (0-336 h) for HUM.

**Table 1:** List of all explanatory variables used in the GLMM analysis. Study areas were ’Zone Armorique’ (ZAM) and ’Vallées et Coteaux de Gascogne’ (VCG).

| Type | Variable | Abbreviation | Unit or coding | Definition |
| --- | --- | --- | --- | --- |
| Spatial & | Study Area | STA | ZAM, VCG | Two study areas |
| Temporal | Year (spring) | YEA | 2012, 2013, 2014 | Three years of sampling |
|  | Week | WEE | #14 – #23 | Week of sampling |
|  | Hour | HOU | h. | Hour of sampling (9am – 8pm) |
| Meteorological | Temperature | TEM | °C | Average temperature over the previous 5 days |
|  | Humidity | HUM | % | Average humidity over the previous 14 days |
| Landscape | Hedgerow perimeter | HEP | m | Sum of perimeters of all hedgerows in a buffer of 350 m |
|  | Woodland perimeter | WOP | m | Sum of perimeters of all woodland plots in a buffer of 350 m |
|  | Water perimeter | WAP | m | Sum of perimeters of all water plots in a buffer of 400 m |
|  | Road perimeter | ROP | m | Sum of perimeters of all roads in a buffer of 400 m |
|  | Building perimeter | BUP | m | Sum of perimeters of all building plots in a buffer of 100 m |
|  | Woodland distance | WOD | m | Minimal distance to the nearest woodland plot |
|  | Water distance | WAD | m | Minimal distance to the nearest water plot |
|  | Road distance | ROD | m | Minimal distance to the nearest road |
|  | Building distance | BUD | m | Minimal distance to the nearest building plot |
|  | Proportion of woodland surface | WOS | % | Proportion of woodlands in a buffer of 350 m |
|  | Surface/perimeter woodland ratio | WOR | m<sup>2</sup>/m | Ratio of woodland surface/perimeter in a buffer of 350 m |
|  | Topography Index | BKT | - | Topography Index (Beven and Kirkby 1979) |

The landscape was characterized in the GIS data by considering seven habitat types: woodland, hedgerow, meadow, crop, road, building, water. Each polygon of the GIS, obtained by human photo-interpretation of the two study areas (ZAM and VCG), was assigned to one of these seven habitats. The distinction between forests (woodlands larger than 50 ha) and woods (woodlands smaller than 50 ha) was not considered inside polygons corresponding to the woodland habitat (see the following parts mentioning the two-part model and the construction of the simulator).

We calculated the perimeter or surface areas of each landscape variable within circular buffers centered on each sampling unit (Table 1). Buffer sizes were determined using circles of increasing diameter, with 10 m steps from 30 to 100 m and 50 m steps from 150 to 400 m. The final selected size was assessed for each variable by the Pearson correlation between nymphal abundance and the given landscape variable. The distance of the sampling unit to the nearest woodland, water, road and building (Table 1) was calculated as described in Figure 2. The topography index of Beven-Kirkby (BKT) (Beven and Kirkby 1979) was calculated from the Digital Elevation Model (DEM) (Zhang and Montgomery 1994) included in the GIS data. The BKT index describes the expected humidity of the soil based on field declivity.

**Figure 2:**
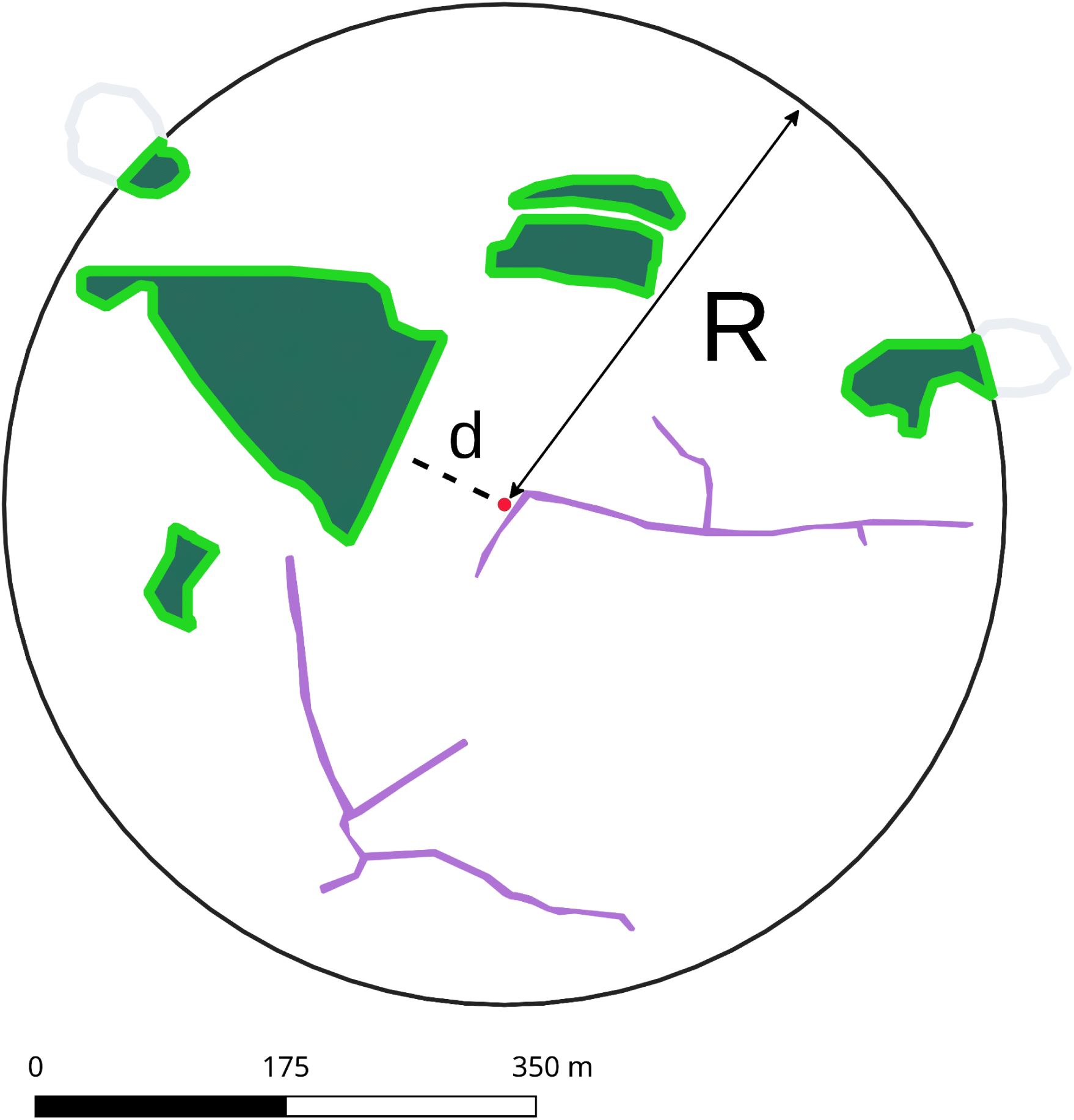
Construction scheme of descriptive landscape variables, illustrated here for the woodland perimeter (WOP). Woodland patches are in green and hedges in purple. The buffer (in black) is centered on one sampling unit (red point). The size of the buffer R is specific to each variable: for WOP, R = 350 m. Around the woodland patch, the perimeter of woodland (in light green) is the sum of perimeters of all woodland patches within the buffer. Distance d (in meter) between the sampling unit here located along a hedge and the nearest woodland patch is indicated by a black dotted line: note that this distance is not limited to the radius of the buffer if the nearest patch is outside the buffer.

### 2.4. Statistical model predicting the abundance of *Ixodes ricinus* nymphs in an agricultural landscape

To model the abundance of *I. ricinus* nymphs, we used a Generalized Linear Mixed Model (GLMM) approach, with a negative binomial distribution (McCullagh and Nelder 1989, Venables and Ripley 2002).

We considered that factors influencing tick density in the forest core were different from those influencing density along hedgerows or woodland borders.Therefore, we built a two-part model, one fitted on the forest core dataset, generalized as “Wood” model, and one applied to the other habitat types, named “Other sector”. Furthermore, hedgerow perimeter (HEP) and woodland distance (WOD) are not relevant for the forest core dataset (see Tables 1 and 2). The predicted number of *I. ricinus* nymphs is then the expectation, *i.e.* the mean. Formally, according to McCullagh and Nelder (1989) and Venables and Ripley (2002), we consider that:

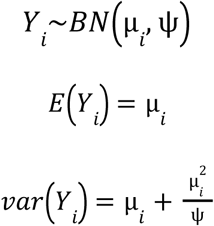

**Table 2:** Results of the GLMM of the explanatory variables for the “Wood” and the “Other sector” models. The explained variable is the abundance of *I. ricinus* nymphs on each sampling unit of 10 m^2^. The observed ranges of the explanatory variables represent the limits later implemented in the simulator; these limits avoid extrapolating simulations beyond those set for the model. The full model with all variables has a DIC of 63.371. Differences of DIC between the full and alternative models are indicated in brackets. Estimates of parameters and precisions are given by the posterior means and standard deviations extracted from MCMC sampling. Parameters that are retained in the models are in bold. The significance of the deviance tests represents the p-values of Chi-square tests: ’***’= 0.001, ’**’ = 0.01, ’*’ = 0.05, ’n.s.’ *>* 0.05. [1] The interaction STA*YEA was preferred over STA + YEA + STA*YEA for parsimony

|  |  |  |  |  |  |
| --- | --- | --- | --- | --- | --- |
| Random effects |  |  |  |  |  |
| STA |  |  |  | *** (+147) |  |
| YEA |  |  |  | *** (+81) |  |
| STA*YEA |  |  |  | *** (+239) [1] |  |
| STA+YEA+STA*YEA |  |  |  | *** (+242) |  |
| Fixed Effects |  |  |  |  |  |
| | | $X^{(W)}$ Wood | | $X^{(O)}$ Other sector | |
|  | Observed range | Test | Estimate | Test | Estimate |
| WEE |  | n.s. | - | n.s. | - |
| HOU |  | n.s. | - | n.s. | - |
| TEM | [7-18] | ** (+8) | **0.07 ± 0.03** | ** (+7) | **0.07 ± 0.02** |
| HUM | [73-89] | n.s. | - | ** (+8) | **0.02 ± 0.004** |
| HEP | [0-7927] | non relevant | - | n.s. | - |
| WOP | [0-10058] | *** (+31) | **2.34 ± 0.35** | *** (+29) | **0.73 ± 0.15** |
| WAP | [0-2326] | n.s. | - | n.s. | - |
| ROP | [0-12454] | n.s. | - | n.s. | - |
| BUP | [0-545] | n.s. | - | *** (+31) | **-2.26 ± 0.27** |
| WOD | [0-621] | non relevant | - | *** (+47) | **-2.62 ± 0.26** |
| WAD | [0-1818] | n.s. | - | n.s. | - |
| ROD | [0-423] | ** (+18) | **1.56 ± 0.34** | n.s. | - |
| BUD | [0-1232] | n.s. | - | n.s. | - |
| WOS | [0-1] | n.s. | - | n.s. | - |
| WOR | [0-1] | n.s. | - | n.s. | - |
| BKT | [0-1] | n.s. | - | n.s. | - |
| Intercept |  |  | **-1.96 ± 1.50** |  |  |

where *Y_i_* is the random variable of the number of nymphs on a sampling unit *i* = 1 *. . . n*. *µ_i_* is the mean of the negative binomial distribution law for the sampling units *i* and *ψ* is the over-dispersion parameter. We assumed that this parameter was common to all sampling units. The GLMM described *µ_i_* in terms of explanatory variables identified as random and fixed effects using a log-link function:

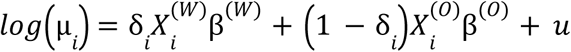

where *δ_i_* is a binary indicator with a value of 1 for the “Wood” model and 0 otherwise. *X_i_^(W)^* is a set of fixed effects associated with parameter *β* ^(W)^, specific to woodlands and *X_i_ ^(O)^* is another set associated with *β*_i_*^(O)^*, specific to the other habitat types. *u* is the set of random Gaussian effects with mean zero and variance matrix *Σ*. For the sake of parsimony, no covariance parameters were included in *Σ*, thus this matrix is diagonal. Variance parameters were estimated in the adjustment process. Random effects were estimated for each sampling unit (among-individual variation) and by group (year of sampling and study area).

To estimate *β*, *ψ* and *u*, we used a Bayesian approach with a Monte-Carlo Markov Chain (MCMC). To identify informative explanatory variables, we first included all the variables in both the sets *X_i_^(W)^* and *β*_i_*^(O)^* to obtain a full model. We identified the informative variables with a backward stepwise selection procedure using the Deviance Information Criterion (DIC), a goodness-of-fit statistic (Spiegelhalter et al. 2002).

### 2.5. MCMC estimation method

The MCMC estimation model is visualised by Directed Acyclic Graph (Spiegelhalter 1998). This graph contains both the likelihood of the data (abundance of *I. ricinus* nymphs), with the *priors* and hyper-*priors* of the informative parameters (Figure 3). The MCMC method used the *posterior* distribution of parameters to simulate samples of *β*, *ψ, u* and the hyper-*priors σ, a* and *b. σ* is a vector of the diagonal of the covariance matrix for random-effects*. a* and *b* are the shape and scale parameter respectively of the Gamma law of over-dispersion. The burn-in was set to 10,000 iterations of the MCMC sample. Sample parameters were constructed by recording one iteration out of 100 for a total of 1,000,000 iterations. Hence, we obtained a total of 10,000 values for each parameter and the posterior means were used as estimations. The choice of *priors* and sample size was based on Robert and Casella (2004) and Robert (2007). The quality of the Markov chains was verified by the Geweke convergence test (Geweke 1992). We used the Stan software (Carpenter et al. 2017) to perform the MCMC simulations.

**Figure 3:**
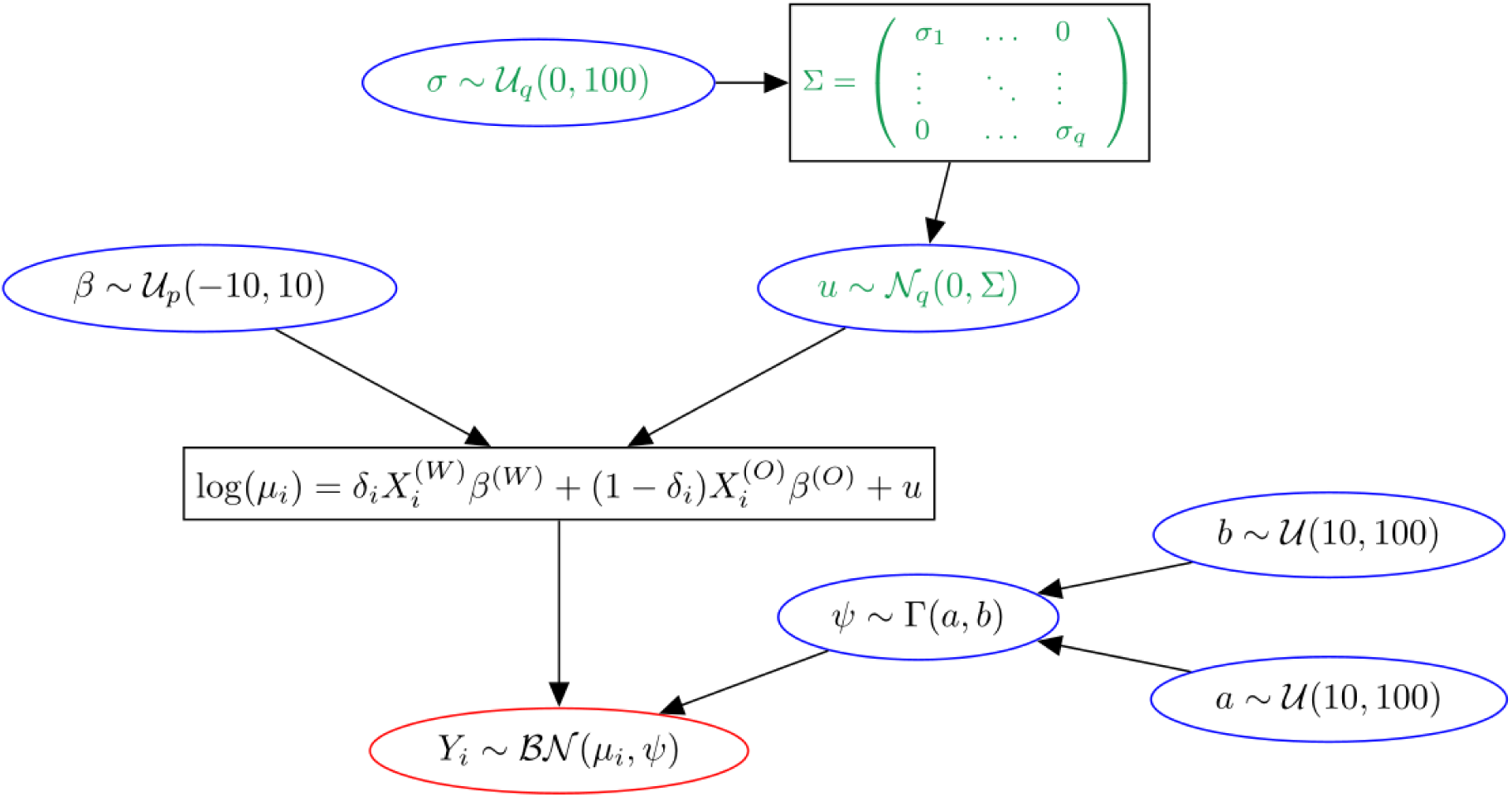
Directed Acyclic Graph of the GLMM model used for the simulator of abundance of *Ixodes ricinus* nymphs *Y_i_*. The ellipses represent random variables while the rectangles correspond to logical nodes. In red, the likelihood of data and in blue the different priors. *µ_i_* is the mean of nymph densities, *ψ* is the over-dispersion parameter. *X_i_* is the matrix of the explanatory variables and *β* in the coefficient vector of the *p* fixed effects. W corresponds to the “Wood” model, and O to the “Other sector” model. *u* is the set of *q* random effects. Σ is a diagonal covariance matrix filled with terms of variance vector *σ*. Γ is the Gamma distribution with parameters *a* and *b*. *U* is the non informative uniform distribution.

### 2.6. Model quality and validation

To check the model quality, we plotted the observed values against the predicted ones without including random effects. We used a bootstrapping method to validate the model on residuals. We split the dataset into 2 subsets. The first, called A, was a random selection of 75% of the whole dataset. The second, B, was the remaining 25%. We used subset A to fit the model parameters. These estimates were then used to perform predictions for the subset B, including random effects. Finally, we computed standardized residuals by plotting observed values against predicted values for the subset B using bootstrapping. We performed 200 iterations of the bootstrapping process to obtain the mean error distribution of the set. For a valid model, a symmetric density curve centered on zero is expected (McCullagh and Nelder, 1989).

### 2.7. Construction of the prototype simulator

To simulate data on a landscape, we used pixels of 30 m x 30 m (which corresponds to CORINE Land Cover resolution) to rasterize the set of habitat polygons with the following rules. When the main habitat of the pixel was defined as woodland (surface of woodland polygons ≥ 75% of the surface of the pixel), the “Wood” model was applied. The other model, named “Other sector”, was applied (1) when a woodland was present but to a lesser extent in the pixel (surface of woodland polygons < 75% of the surface of the pixel) or (2) when hedgerows were present at the center of the pixel. In order to allow hedgerows to be considered in some pixels, which would otherwise be almost impossible due to their small size, a 7.5 m wide spatial buffer was added by the simulator to the surface around the hedgerow polygons. This “Other sector” model corresponds thus to (1) woodland borders or to (2) hedgerows (Figure 4). As *I. ricinus* abundance is known to drastically fall a few meters away from a woodland or a hedgerow (Medlock et al. 2020), we did not predict tick abundance outside those two habitats (*i.e.* in meadows, crops, roads, buildings and water): thus, when the conditions described above were not met in a pixel, no model was applied and the abundance of nymphs was set to “not simulated” (Figure 5). A prototype simulator named ‘OSCAR: tick abundance simulator’ was built as a *R shiny* application (https://oscar-abondance.sk8.inrae.fr/). The calculation part of the simulator was written in C++ using the GDAL library, which allows parallelization of geo-processing of raster and vector data (Rouault et al. 2025). The *R shiny* part is the web user interface of the simulator hosted by the SK8 project of INRAE (https://sk8.inrae.fr/).

**Figure 4:**
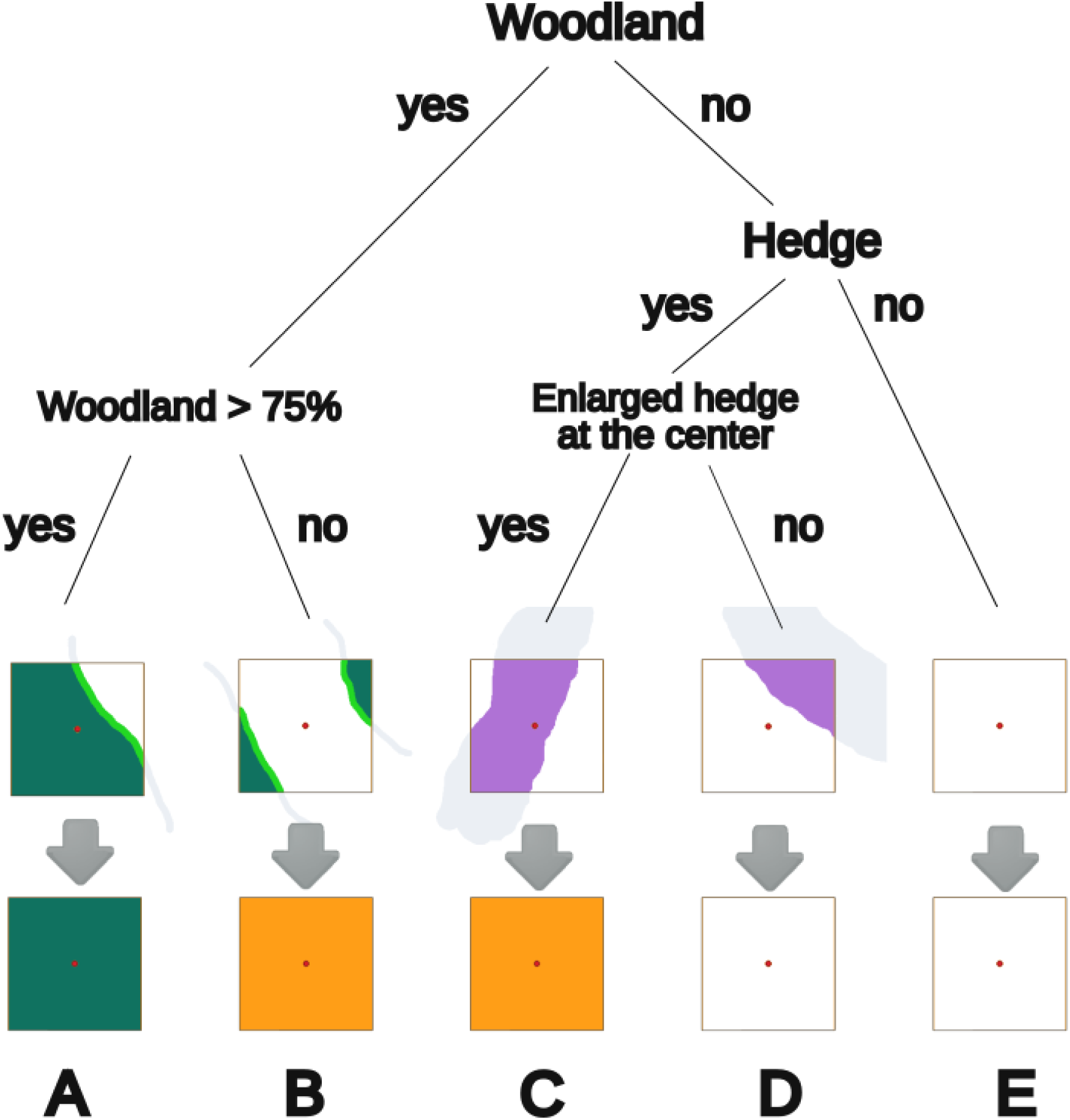
Decision tree implemented in the simulator to classify the 30 x 30 m pixels as “Wood”, “Other sector” or “Not simulated”. Each class leads to the corresponding model. A) presence of a woodland habitat with a wooded area covering more than 75% of the surface of the pixel (*i.e.* more than 75% of 900 m²): “wood” class. B) presence of a woodland habitat with a wooded area covering less than 75% of the surface of the pixel: “Other sector” class. C) absence of a woodland habitat, presence of a hedge and, after its enlargement (7.5 m wide spatial buffer), presence of the hedge at the center of the pixel: “Other sector” class. D) absence of a woodland habitat, presence of a hedge and, after its enlargement, absence of the hedge at the center of the pixel: “Not simulated” class. E) absence of a woodland habitat and absence of a hedge: “Not simulated” class.

**Figure 5:**
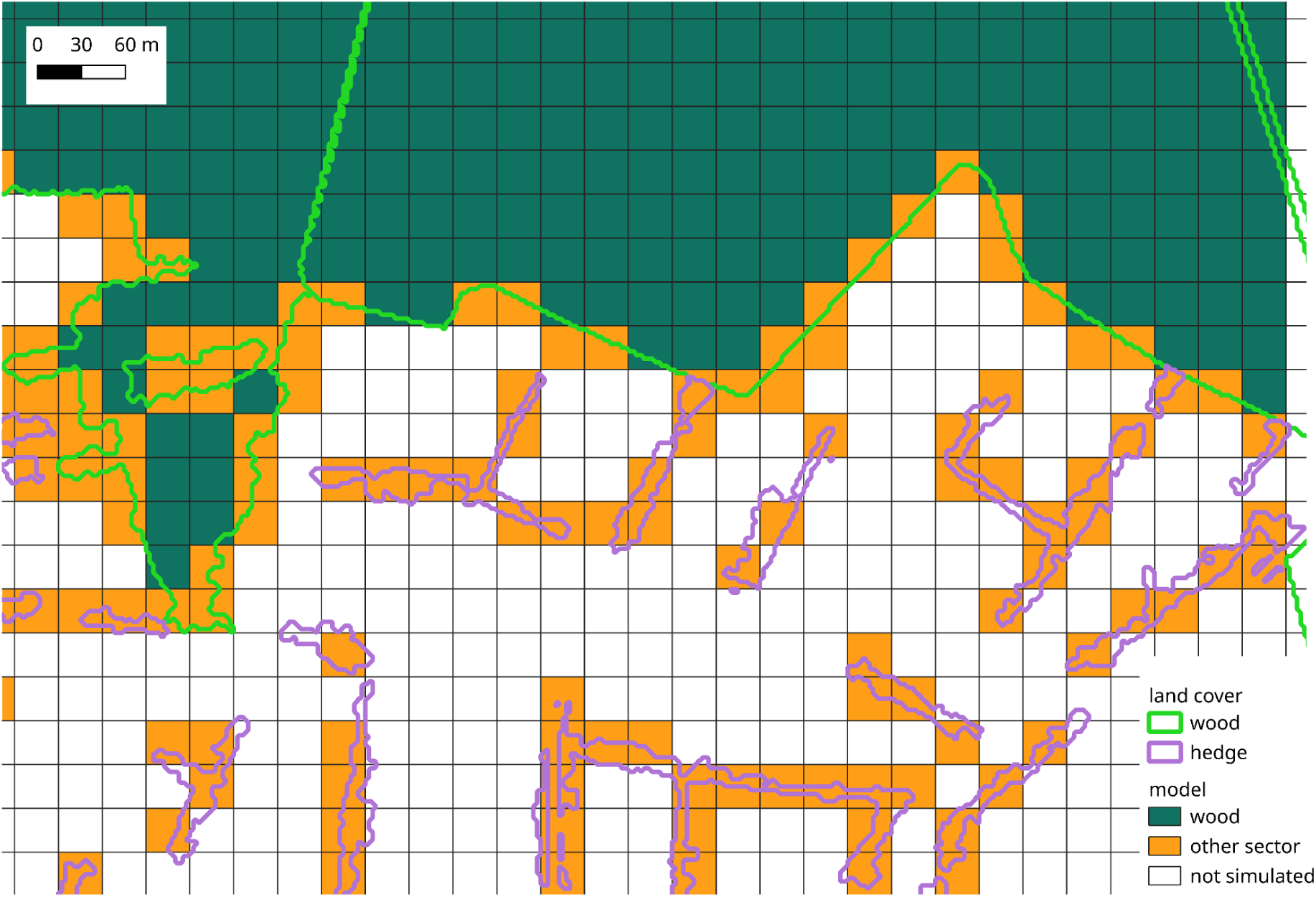
Example of the class of each pixel of a landscape obtained after the application of the decision tree (Figure 4). Each 30 x 30 m pixel is classified as “Wood”, “Other sector” or “Not simulated”.

In order to be usable for new landscapes by future users of the simulator, the landscape input data must be in the form of GIS files (vectorial ESRI shapefile format), with landscape descriptors corresponding to one polygon per patch of habitat. The landscape descriptors have to include polygons of woodlands, hedgerows, meadows, crops, roads, buildings and water. The meteorological parameters, *i.e.* temperature and humidity, were fixed in the simulator to the means of the data used for the estimation of the parameters of the model. The abundance of *I. ricinus* nymphs is standardized, becoming unitless, and represented on a map with a color gradient from 0 (min) to 1 (max). This raster can be downloaded in GeoTiff format.

## 3. Results

### 3.1. Overview of field data

In total, over the three years, 11,998 nymphs of *I. ricinus* were collected on the vegetation of the 5,390 sampling units, representing a mean abundance of 2.23 nymphs per 10 m^2^ (min = 0; max = 74; median = 1). The 90 geo-referenced transect lines on each area are represented in Figure 6, with their average nymphal density calculated over the three years per 100 m^2^. Details on the dataset are given in the data paper by Lebert et al. 2020.

**Figure 6:**
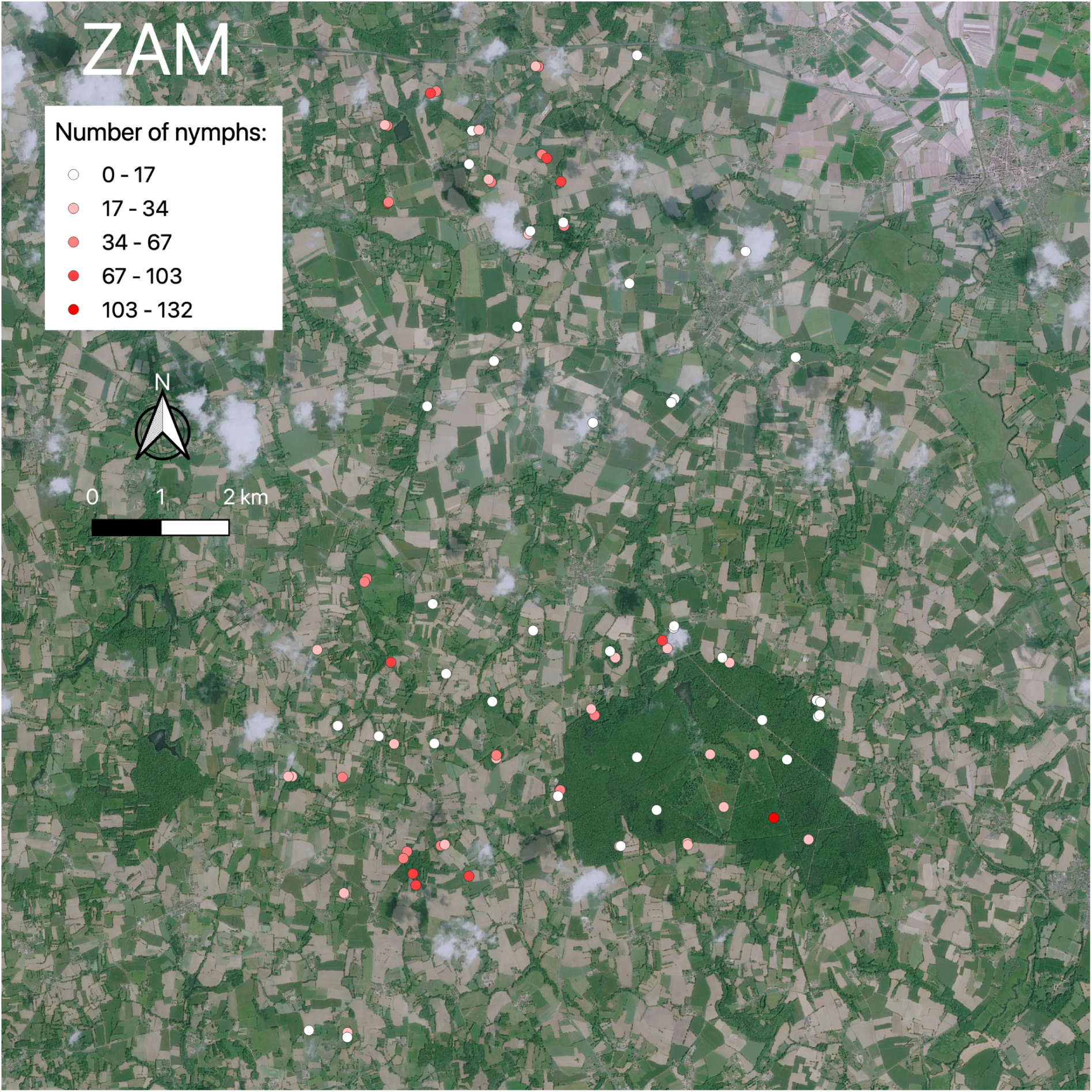
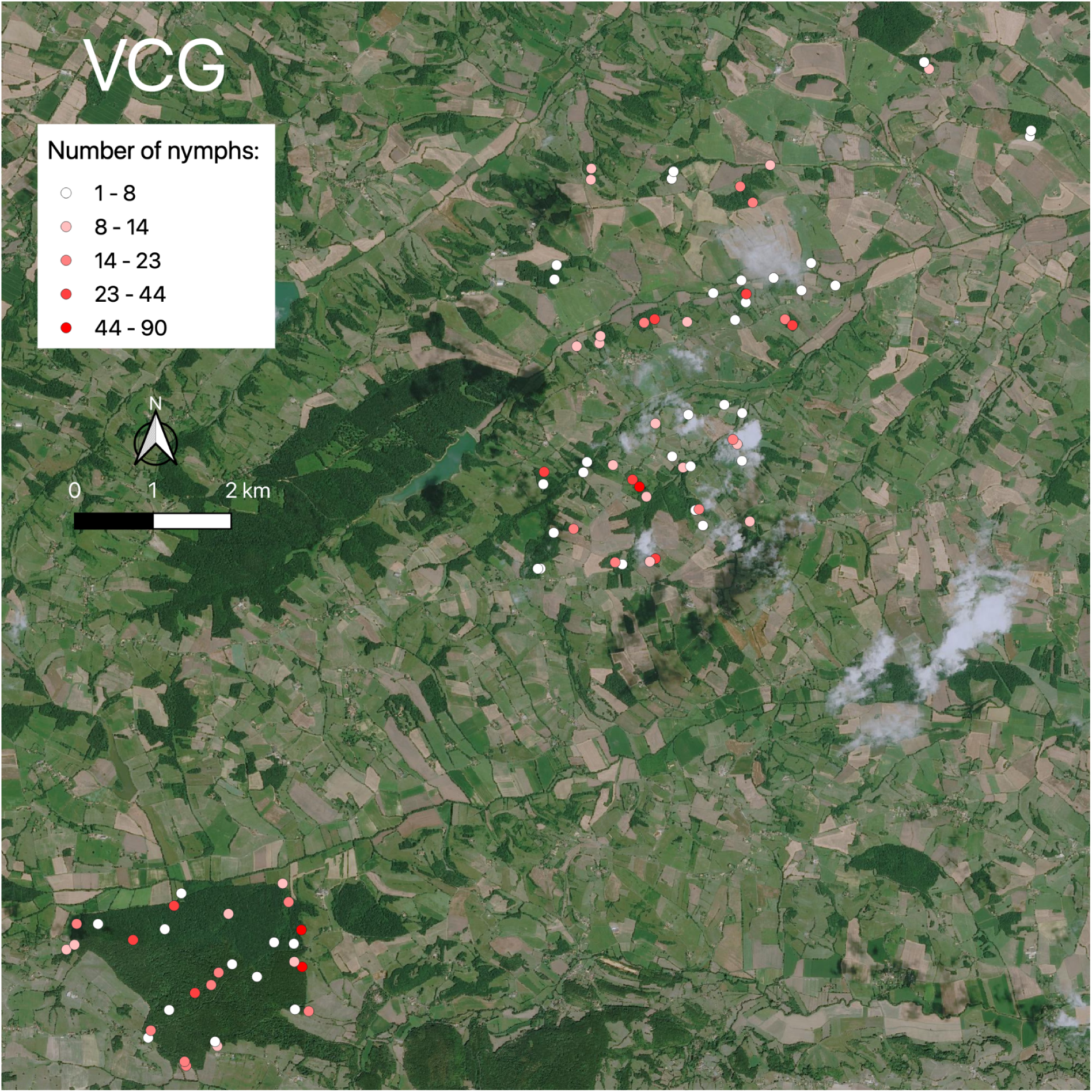
Localization of the 90 line-transect centroids (represented as spots) on each study area (ZAM, VCG), with their average nymphal density calculated in spring over the three years. Average nymphal density at the line-transect scale ranged from 0 (white spot) to 132 individuals (darkest red spot) in ZAM (median=23; mean=31), and from 1 to 90 in VCG (median=10; mean=14). Each transect line corresponded to 10 sampling units, *i.e.* 100 m^2^. Background map from SPOT 6 satellite.

### 3.2. GLMM results, predictive equation, model quality and validation

Results of the GLMM model are presented in Table 2. All the random effects were informative, but for parsimony we selected only the interaction between study area and year (STA*YEA). Neither the fixed effect of week number nor the hour of sampling were retained for either of the two models (“Wood” and “Other sector”). For the “Wood” model, temperature (TEM), woodland perimeter (WOP) and road distance (ROD) were positively linked to nymphal abundance. For the “Other sector” model, nymphal abundance was positively linked to both temperature (TEM) and humidity (HUM) but also to woodland perimeter (WOP), and negatively linked to woodland distance (WOD) and building perimeter (BUP).

Based on the consideration of informative environmental variables (Table 2), the predicted number of *I. ricinus* mean nymphs quantity was:

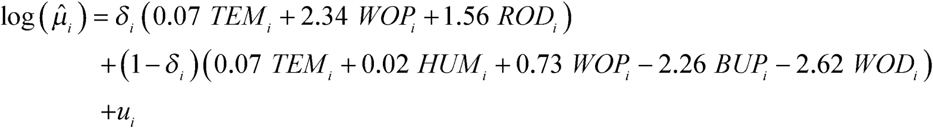

The over-dispersion parameter *ψ* was estimated as 0.33 *±* 0.03. It is thus possible to obtain the simulated number of nymphs at sampling unit *i* using the distribution law:

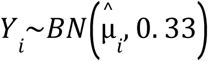

The quality plot (Figure 7) shows a suitable fit between the predicted and observed values, except for the few observations with high density values. The validation plot (Figure 8) illustrates that the density of mean simulation errors is globally centered around 0 and symmetrical.

**Figure 7:**
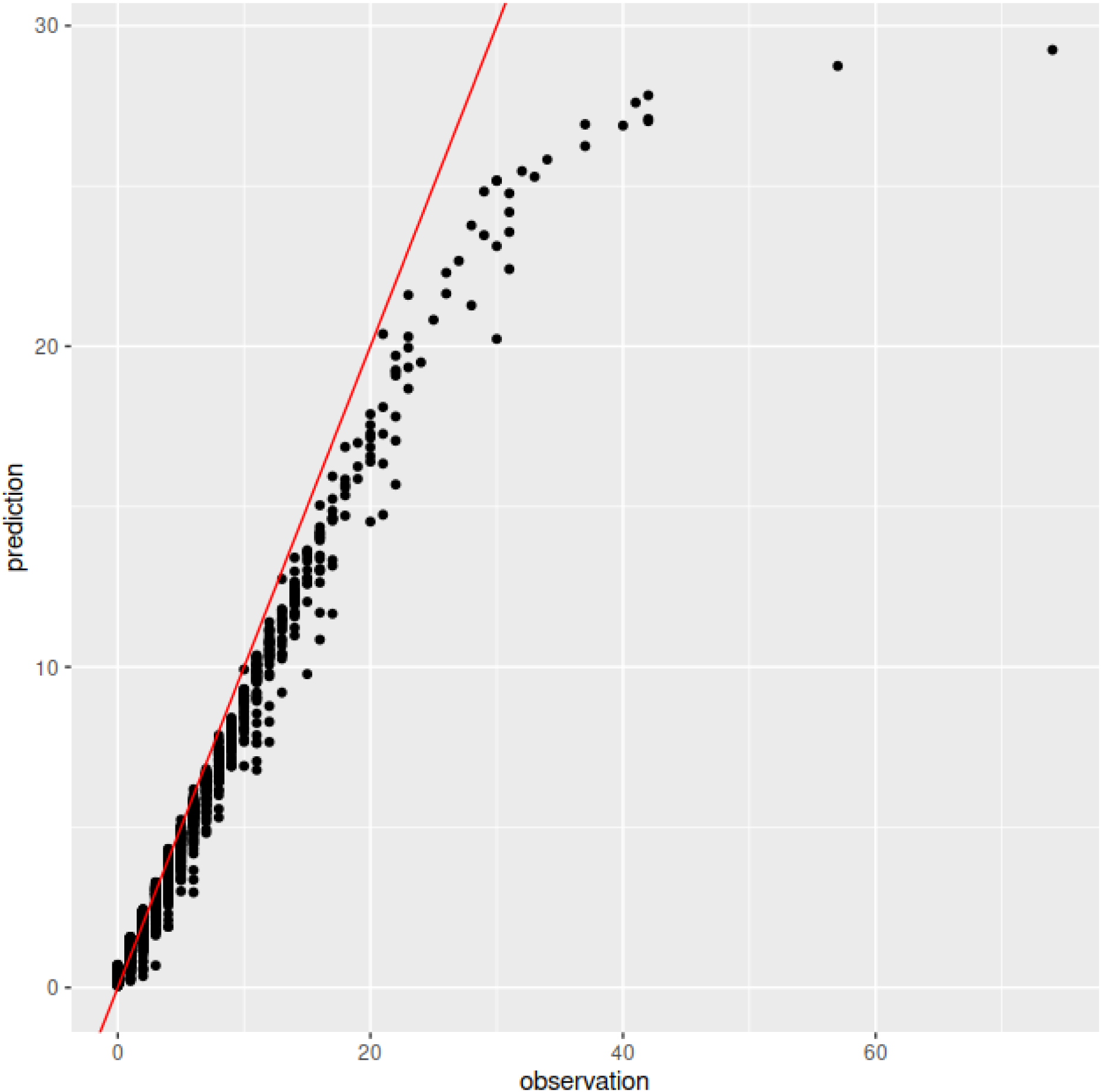
Observed *versus* predicted number of nymphs on 10 m^2^ for the 5390 sampling units. The first bisectrix (red line) is indicated here as a reference.

**Figure 8:**
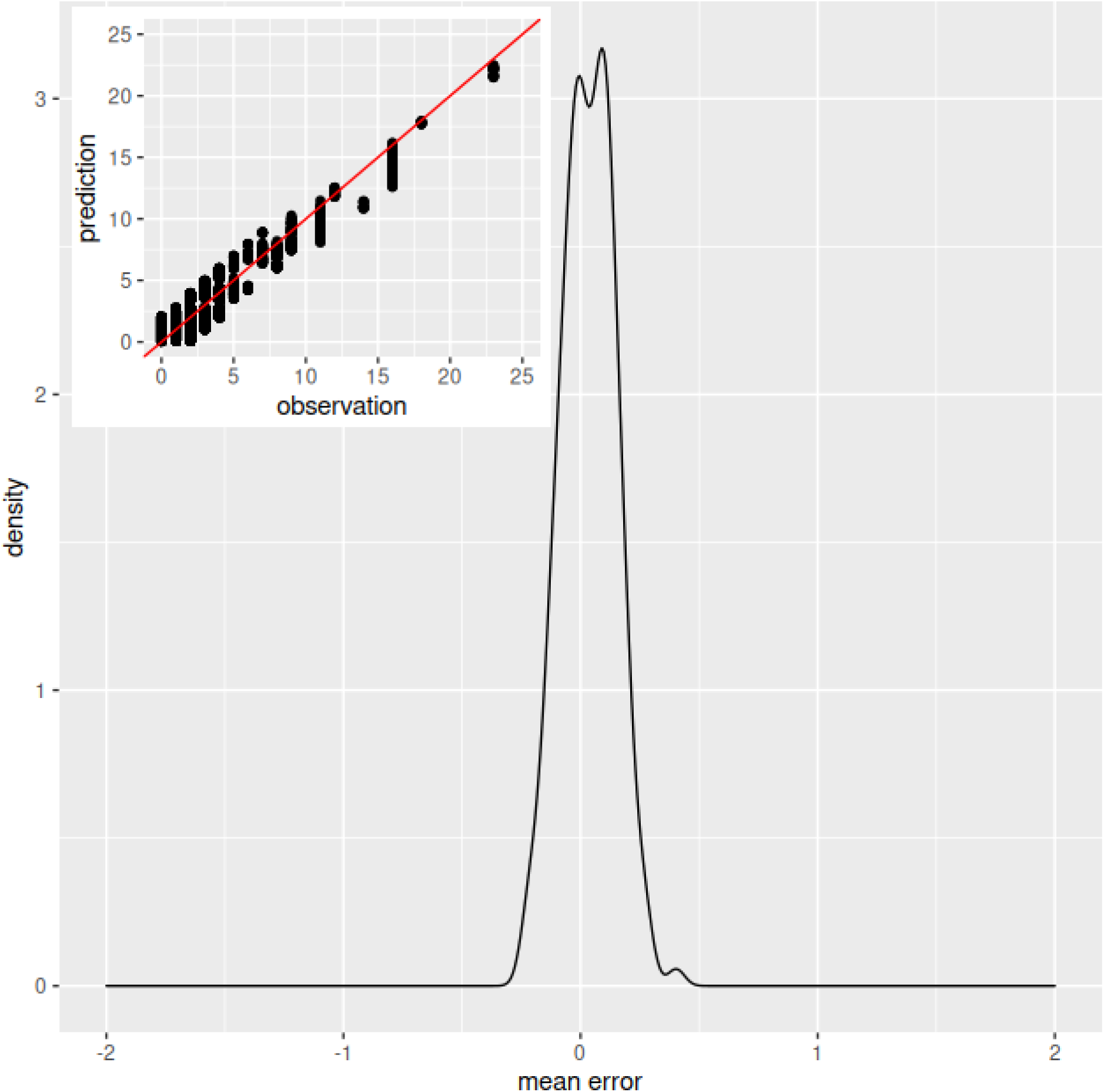
Validation process using one quarter of the collected data that was not used in parameter estimation (n = 1347 sampling units). A total of 200 simulations were run using the estimated parameters. A scatter plot of the predicted *versus* observed values for one of the 200 simulations is presented in the upper left corner..

### 3.3. Simulated nymphal abundance at the landscape scale

Figure 9 presents the application interface of the simulator. In this figure, the coding convention (A), the main landscape characteristics (D) and the simulation information (G) are presented. The user’s graphical control elements for input are also presented (B, C, E and F). Figure 10 gives an example of the simulation output. Landscape and nymphal simulated abundance are plotted at the same spatial scale and the user changes both scales simultaneously on the simulator. Within woodland pixels, where the “Wood” model applies, the positive effect of distance to the nearest road (ROD, see Table 2) on tick abundance can be easily visualized. The positive effect of woodland perimeter (WOP) is less conspicuous, but more noticeable in small woodland patches ; in central locations within woodlands, without roads or clearings in the vicinity, woodland perimeter is invariable, as it corresponds to the perimeter of the circular buffer: thus, in these locations, nymphal densities tend to be high and constant. In the pixels where the “Other sector” model applies (*i.e.* woodland borders and hedgerows), the positive effect of woodland perimeter (WOP) and the negative effects of woodland distance (WOD) and building perimeter (BUP) are more difficult to visualize due to interactions among those variables in such heterogeneous landscapes.

**Figure 9:**
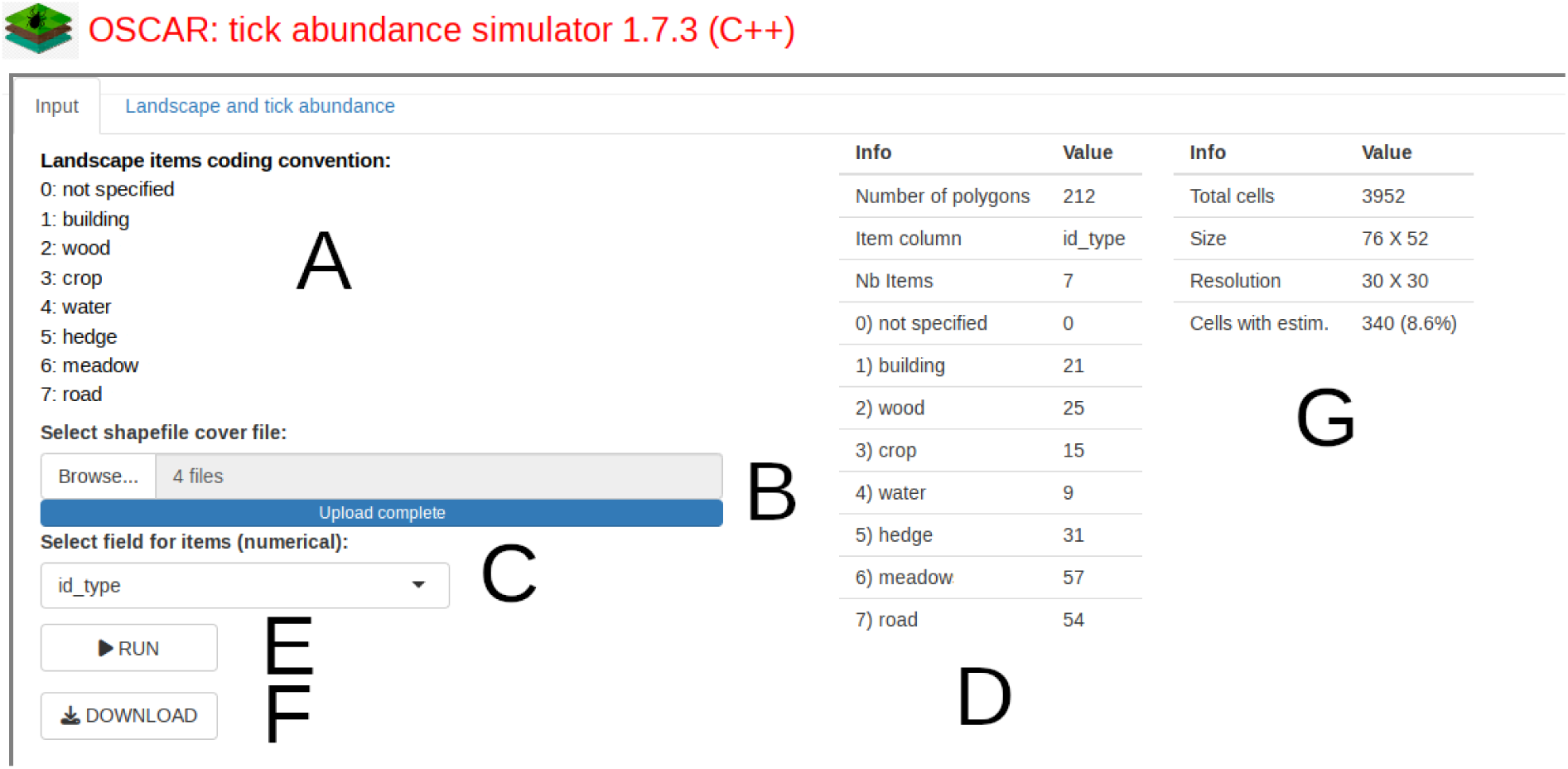
Input page of the “OSCAR: tick abundance simulator” web interface. A) List of the landscape items. B) Selector of vector data files used as input data to describe the landscape (ESRI shapefile format). C) Selection of the column (field) of attribute data corresponding to coding landscape items. D) Number of polygons for each landscape item. E) Run command to compute simulation. F) Download command of the raster output file. G) Information about the simulation process (total number of cells, size of the cell - 30x30 m - and number of cells with tick density estimates).

**Figure 10:**
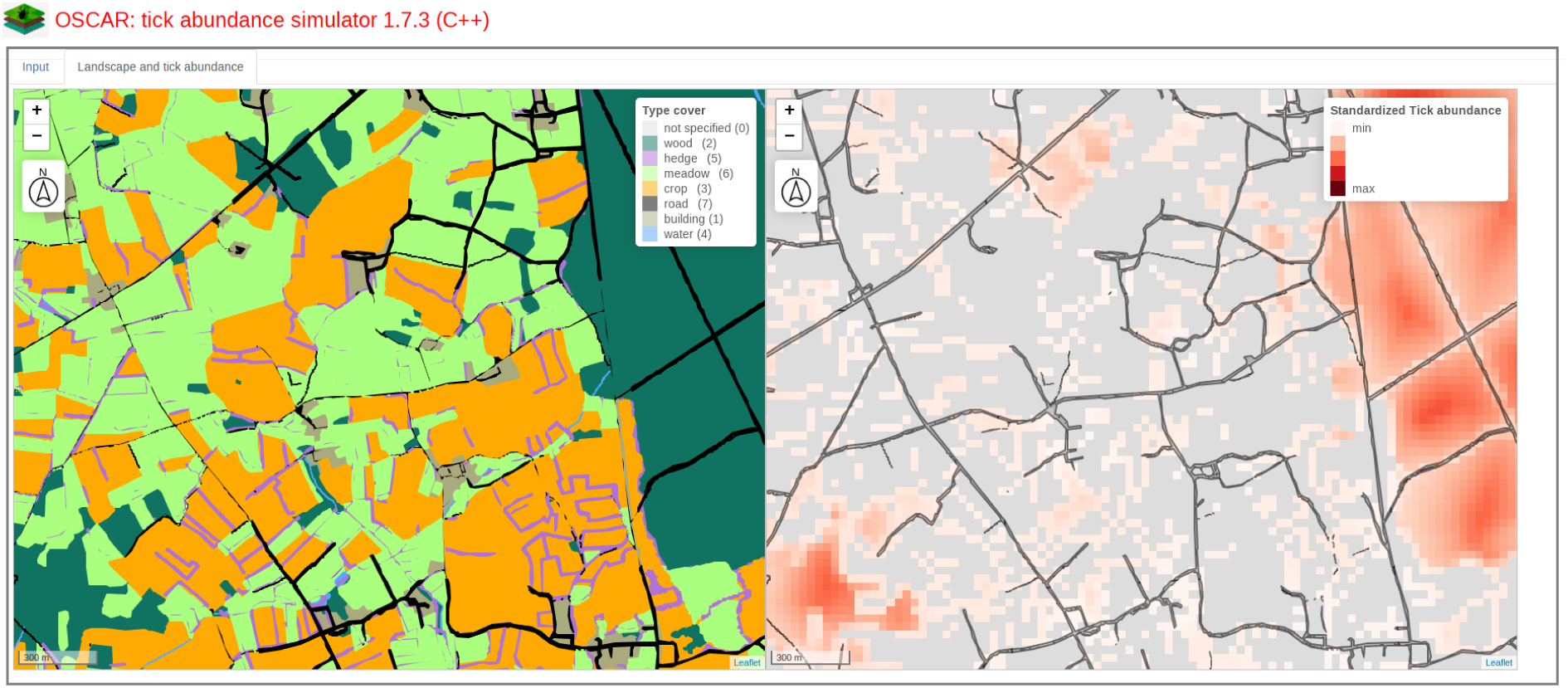
Output page of the “OSCAR: tick abundance simulator” web interface (part of ZAM). On the left, the input vector data used for simulation. Each patch is a polygon with a landscape item numbered from 1 to 7 (0 corresponding to “not specified”). On the right, the tick abundance raster is visualized at a standardized scale with the darkest red pixels corresponding to the highest tick density. Grey pixels correspond to areas where tick density is not simulated (*i.e.* outside cells corresponding either to woodlands or to hedgerows). Roads are represented to facilitate geographical location.

## 4. Discussion

The aim of this study was to model the spatial heterogeneity in tick abundance using an extensive data set from an observational study in two pilot areas, taking into account landscape features at a fine spatial grain, to identify the main drivers of this heterogeneity. Subsequently, the identified environmental drivers were used to simulate spatial variation in tick abundance at the landscape scale. This simulator can be used on any map provided by the user to visualize the expected spatial heterogeneity of tick density.

### Identifying the most relevant drivers explaining tick abundance at the landscape scale

The prototype simulator is based on a statistical model that provided a satisfactory fit with the field data (Figure 7). These data were collected with the aim of maximizing signal and accuracy in terms of nymphal tick abundance: the sampling protocol was performed during the spring peak of nymphs for three consecutive years on two study areas rich in woodlands, and therefore *a priori* favorable to ticks. This extensive dataset therefore allowed us to make a thorough analysis of the link between tick densities during their peak activity (spring) and landscape characteristics in fragmented agroecosystems.

Results from the GLMM modeling highlight the influence of both abiotic and biotic factors on spring tick densities; the driving factors explaining tick abundance differed depending on whether ticks were collected inside or outside woodlands. Among abiotic variables, temperature was found to influence tick density both inside and outside woodlands, whereas humidity was only important outside. The positive effect of spring temperature on tick density is in agreement with regard to its well-known functional role in tick development (Randolph et al. 2002) and activity (Perret et al. 2003, Vail and Smith 1998). However, tick activity and survival are also considered to strongly depend on relative humidity (Daniel et al. 1976). Relative humidity may not be a limiting factor inside woodlands, as observed in the present study, due to (i) the shade provided by the canopy, (ii) a reduction in wind velocity by the vegetation of different heights and (iii) the buffering effect of the leaf litter.

Regarding landscape characteristics, the length of woodland perimeter around the investigated sampling point (WOP, in a buffer of 350 m) was found to positively influence tick densities. This effect is mostly observed inside woodlands, but also outside (Table 2). The length of the woodland perimeter is a function of both woodland occurrence and the interface between woodlands and surrounding habitats. The positive effect of the length of woodland perimeter could be related to higher abundance of tick hosts (e.g. small mammals and roe deer) in these ecotones; indeed, many of these hosts use more open areas for feeding and woodland habitats for shelter (Bonnot et al. 2013, Rigoudy et al. 2024), favoring tick occurrence.

Within woodlands, road distance (ROD) is positively linked to tick abundance, such that tick abundance increases as the distance to the nearest road increases. This may be due to a repulsive effect of roads on hosts. Such an effect has already been reported for roe deer in several studies (Bonnot et al. 2013).

Outside woodlands, the distance to the nearest woodland (WOD) is negatively linked to tick abundance. This is in accordance with Medlock et al. (2020), who found higher *I. ricinus* nymphal densities along woodlands than along hedgerows, located further away from woodlands. This may be linked to the fact that woodlands provide the most suitable habitat for tick survival (notably due to higher humidity) and could be considered as sources of ticks for other suitable habitats, such as hedgerows, as suggested by Hoch et al. (2010) in a modelling study. The effect of woodland distance on tick abundance could be due to the influence of woodland on roe deer habitat use, more abundant closer to woodlands, where they can find refugia against human disturbance (Benhaiem et al. 2008, Martin et al. 2018, Morellet et al. 2011, Padié et al. 2015).

Finally, tick densities were higher outside woodlands when there were less buildings in the surroundings (negative effect of building perimeter (BUP) in a buffer of 100 m), which might be related to avoidance of proximity with human presence by roe deer (Bonnot et al. 2013, Coulon et al. 2008), as mentioned above for the effect of road distance (ROD).

### Simulating tick abundance in landscapes

Subsequently, we used the environmental drivers identified as informative in our statistical model to simulate spatial variation of tick abundance in relation to landscape structure. Simulated nymphal abundance, and hence related risk, appeared to be maximal in large forests, especially in areas situated far from roads. However, some isolated woodlands harbored a high nymphal abundance due to the positive effect of the length of woodland perimeter.

This article illustrates the feasibility of simulating tick abundance at the scale of an heterogeneous agricultural landscape. Any user who has access to a map with the appropriate environmental features in such a landscape would then be able to generate an output map of predicted tick abundance. Validation of the simulator and its underlying statistical model in new areas would increase generalizability for wider application. This work represents therefore the first step towards providing a tool that would help stakeholders to manage tick-borne risk in agricultural habitats.

A perspective would consist in coupling the statistical model of nymph abundance with a theoretical landscape model (Papaïx et al. 2014, Poggi et al. 2018). Such a coupling would allow the analysis of the effect of landscape characteristics (*i.e.* fragmentation and connectivity, defined on simulated landscapes) on the risk associated with ticks, and therefore give insight into landscape management measures that could help to lower these risks.

## Acknowledgments

We acknowledge the directors of the two ‘Zone Atelier”, Zone Atelier Armorique’ (https://osur.univ-rennes1.fr/zaar-home) and PYGAR, https://pygar.omp.eu/ for allowing us to conduct our investigations in those studies. We thank all the colleagues involved in the OSCAR project listed in the data paper Lebert et al. 2020 for their help for data acquisition. We thank Patrick Gasqui (INRAE, UMR EPIA) and Séverine Bord (INRAE, UMR LISIS) for statistical advice and Jean-François Rey(INRAE, CATI IMOTEP) for his help to use R-Shiny facilities (Project SK8). We thank the TMT (‘Tiques et maladies à Tiques’) currently linked to the SFE2 for constructive reflections and constructive ideas when preparing this manuscript.

